# Behavior of cetaceans in the waters south of Pico Island (Azores, Portugal) - data obtained during whale-watching 2020

**DOI:** 10.1101/2022.05.29.493883

**Authors:** Peter Zahn, Lynn Kulike, Armin Bloechl

**Author notes:** Corresponding author: Peter Zahn.

## Abstract

Land-based and boat-based surveys were conducted to collect data during whale-watching excursions in July/August and October 2020. Occurrence and behavior of cetaceans south of Pico Island (Azores) were determined. 10 species were sighted. The most frequently sighted species were: Sperm whale (*Physeter macrocephalus*), Sei whale (*Balaenoptera borealis*), Atlantic spotted dolphin (*Stenella frontalis*), Common bottlenose dolphin (*Tursiops truncatus*), and Risso’s dolphin (*Grampus griseus*). 51 different behaviour patterns were recorded. The majority, 32 behavior, were observed from two or more species. 19 behavioral patterns were observed only once from a single species, Atlantic spotted dolphins showed nine and Sperm whale showed six. The most behavioral patterns were displayed by *Stenella frontalis* with 26 and *Pseudorca crassidens* with 22. Interspecific cooperative hunting and intraspecific food sharing were observed, for the latter there were no known observations in the Azores so far.

## Introduction

Many people are highly attracted by cetaceans. They approach boats, ride in the bow wave and perform astonishing acrobatics, apparently of pure pleasure. Therefore, many are interested in their behavior. The Greeks already reported bow riding in the Mediterranean Sea by likely Common bottlenose dolphins, Shortbeaked common dolphins, and Striped dolphins (Würsig et al, 2018). Behavior can sometimes be useful for identification purposes, because it varies greatly between species (Carwardine, 2020; Still et al, 2019). In this study, behavior was used, as a first aim to assist in the identification of the species. The second aim was to investigate of how many different behavior patterns were possible to observe.

Places that were once popular for whaling are nowadays often used for whale-watching. Tourism is growing rapidly and also whale-watching activities (Visser et al, 2011b). The Azores archipelago developed into a hotspot of whale-watching since the early 1990s. Cetaceans are an important marine megafauna group in the Azores, with 28 species recorded so far (Silva et al., 2014), and are probably a key component of the Azores marine ecosystem (Tobeña et al., 2016). Information on behavioral patterns of cetaceans in this region remains limited (Cechetti et al 2018). Surveys are costly, and surveying large areas of offshore waters comprise a lot of logistic and operational difficulties. The use of other data sets constitutes a valuable alternative for investigating how cetaceans use these areas (Silva et al, 2014). Whale-watching activities offer a source of valuable data and funding for cetacean research (Pereira, 2008a; Steiner et al., 2012).

Some limitations, e.g. that the study area may not be equally covered, or that the findings of this research may not be representative for the entire research area, need to be discussed. Nevertheless, the data collected on marine life as a “by-product” of whale-watching may contribute crucial data for understanding occurrence of cetaceans around the island of Pico (Bron et al, 2019). Precise knowledge of the distribution and behavior of the species is important for efficient protection of whales. Whale-watching tours are offered possibly year-round. Therefore, it is a potential tool for detecting long term changes. The excursion aims to gain some data of the behavior of the whale species observed in the waters of the south coast of Pico Island.

## Material and Method

In this study, data were collected in the Azores, an archipelago composed of nine volcanic islands in the North Atlantic Ocean, during one land-based and 20 boat-based surveys in the summer and autumn months of 2020. 7 days in July, 2 in August and 5 days in October. They were collected during commercial trips with the whale-watching company Espaço Talassa, with its homeport in Lajes do Pico. Whale-watching companies rely on land-based lookouts to detect cetaceans. They are usually located in a fixed land station. These “vigias” directed the vessels to the whales and dolphins.

The vantage point of Espaço Talassa is located on the south coast of Pico Island, near the town of Lajes. The “vigia” is about 200 m away from the cliff edge, at an altitude of 75 m, and with the approximate position of 38° 23’ 094” N and 28° 14’ 498” W. Using 15x or 20x binoculars, occurrence and movements of cetacean groups were recorded. The lookout may see up to 35 km in perfect weather conditions. Based on the 210° degree of a circle centred on the observation point, the observation field amounts to 1900 km^2^ (Pereira, 2008b). The “vigia” was in permanent radio contact with the boat, the crew confirmed the identification.

The excursion consisted of a group of 12 people in altogether, students and lecturers. The boat surveys were led by two skippers that represents the maximum capacity of 14 participants for one whale-watching boat. From this it follows that the number of trained observers were at least three. Each boat tour took three hours and occurred in the morning or in the afternoon respectively. A de-briefing after every boat-tour was carried out. All sightings were reviewed and questions answered. A comparison with the notes made on board was performed. Data collection was restricted. No tracked sail routes. Recorded number of trips 20. Rough indicator of effort by determining the number of species encountered per trip.

## Results

During the study, a total of 10 whale species were observed. Representatives of four families were recorded. One species from each of the three families Balaenopteridae, Physeteridae and Ziphiidae were sighted. With six genera and seven species, the representatives of the family Delphinidae were seen most frequently. In relation to their feeding ecology, two main groups are distinguished. Four teutophagous and deep diving species, *Physeter macrocephalus, Mesoplodon bidens, Globicephala macrorhynchus*, and *Grampus griseus*. The other six species feed on fish, squid or crustaceans, *Balaenoptera borealis, Delphinus delphis, Pseudorca crassidens, Stenella coeruleoalba, Stenella frontalis*, and *Tursiops truncatus* (Carwardine, 2020).

Table 1 shows the number of sightings and the number of behavioral patterns recorded by each cetacean species observed. *Physeter macrocephalus, Balaenoptera borealis*, and *Stenella frontalis* were the most sighted species. Atlantic spotted dolphin, False killer whale, Risso’s dolphin, Common bottlenose dolphin, and Sperm whale displayed the most behavior patterns. All Mesoplodon species were too far away and further careful approaches failed and no behavior was recorded. The field data compiled during the survey encompass 51 different behavior patterns.

**Tab.1:** Observing behavior of sighted whale species (Behnsen; Freitag; Marks; Sordyl; Zahn).

| Species | Behavior (July 24<sup>th</sup> to August 2<sup>nd</sup> 2020) | Behavior (October 6<sup>th</sup> to 10<sup>th</sup> 2020) | No. of sightings | No. of behavior |
| --- | --- | --- | --- | --- |
| ***Balaenoptera borealis***<br>(Sei whale) | Aggregation (with *Stenella frontalis*)<br>Blowing<br>Breaching<br>Diving<br>Feeding<br>Surfacing<br>Swimming lateral<br>Traveling | Approach boat<br>Blowing<br>Curiosity<br>Diving<br>Feeding<br>Surfacing<br>Traveling | 20 | 10 |
| ***Physeter macrocephalus***<br>(Sperm whale) | Aggregation (*Delphinus delphis*)<br>Approach boat<br>Blowing<br>Breaching<br>Curiosity (dive on the boat)<br>Diving<br>Fluking<br>Lob-tailing<br>Pre-dive flex<br>Resting<br>Surfacing<br>Traveling slow<br>Traveling | Approach boat<br>Blowing<br>Curiosity<br>Display (male towards boat)<br>Diving<br>Fluking<br>Joining (mother and calf)<br>Lactating<br>Logging<br>Play (Juvenile)<br>Pre-dive flex<br>Resting<br>Shallow Dive<br>Surfacing<br>Swimming lateral | 30 | 20 |
| ***Mesoplodon bidens***<br>(Sowerby's Beaked whale) | Resting |  | 1 | 1 |
| ***Delphinus delphis***<br>(Short-beaked Common dolphin) | Aggregation (*Physeter macrocephalus*)<br>Approach boat<br>Association (*Stenella coeruleoalba*)<br>Blowing<br>Bow riding<br>Interspecific cooperative hunting (*Stenella coeruleoalba*)<br>Diving<br>Feeding<br>Fluking<br>Leaping<br>Surfacing<br>Traveling<br>Traveling fast |  | 6 | 13 |
| ***Globicephala macrorhynchus***<br>(Short-finned Pilot whale) | Blowing<br>Diving<br>Peduncle arch<br>Resting<br>Slapping tail<br>Surfacing<br>Synchronous behavior<br>Traveling |  | 8 | 8 |
| ***Grampus griseus***<br>(Risso's dolphin) | Aggressive behavior (*Tursiops truncatus*)<br>Association (*Tursiops truncatus*)<br>Blowing<br>Communication (adults joining)<br>Diving<br>Drowning (*Tursiops truncatus*)<br>Fluking<br>Joining (adult males join juveniles)<br>Lateral swimming<br>Leaping<br>Slapping flipper<br>Slapping tail<br>Socializing<br>Surfacing<br>Synchronized behavior<br>Traveling slow<br>Traveling<br>Traveling fast | Blowing<br>Diving<br>Fluking<br>Slapping flipper<br>Slapping tail<br>Socializing<br>Sprint (40 m)<br>Surfacing<br>Surfing<br>Traveling | 12 | 20 |
| ***Pseudorca crassidens***<br>(False killer whale) |  | Approach boat<br>Association (*Tursiops truncatus*)<br>Blowing<br>Bow riding<br>Breaching<br>Diving<br>Feeding<br>Fluking<br>Food sharing<br>Hunting<br>Hunting cooperative interspecific (*Tursiops truncatus*)<br>Lining (> 1 km)<br>Lunging<br>Slapping flipper<br>Slapping tail<br>Socializing<br>Spy-hopping<br>Surfacing<br>Swimming lateral<br>Synchronous behaviour<br>Traveling<br>Traveling fast | 4 | 22 |
| ***Stenella coeruleoalba***<br>(Striped dolphin) | Association (*Delphinus delphis*)<br>Blowing<br>Diving<br>Hunting cooperative interspecific (*Delphinus delphis*)<br>Leaping<br>Porpoising<br>Surfacing<br>Surfing |  | 4 | 8 |
| ***Stenella frontalis***<br>(Atlantic Spotted Dolphin) | Aggregation (*Balaenoptera borealis*)<br>Approach boat<br>Blowing<br>Bow riding<br>Diving<br>Feeding<br>Hunting<br>Leaping<br>Leaping (acrobatic)<br>Lunging<br>Mating<br>Snout riding (*Balaenoptera borealis*)<br>Splashing (all somersaults)<br>Surfacing<br>Traveling | Approach boat<br>Association forming superpod<br>Association (*Tursiops truncatus*)<br>Blowing<br>Bow riding<br>Bubble blowing<br>Diving<br>Feeding<br>Fluking<br>Hunting<br>Hunting cooperative interspecific (*Tursiops truncatus*)<br>Leaping<br>Lining<br>Pursuit<br>Slapping head<br>Slapping tail<br>Slapping tail to stun fish<br>Sound vocalization<br>Surfacing<br>Surfing<br>Tail-Standing<br>Traveling | 17 | 26 |
| ***Tursiops truncatus***<br>(Common bottlenose dolphin) | Approach boat<br>Aggressive behavior (*Grampus griseus*)<br>Association (*Grampus griseus*)<br>Blowing<br>Bow riding<br>Diving<br>Drowning (*Grampus griseus*)<br>Fluking<br>Leaping<br>Sprinting<br>Spy-hopping<br>Surfacing<br>Traveling | Approach boat<br>Association (*Pseudorca crassidens*)<br>Association (*Stenella frontalis*)<br>Blowing<br>Bow riding<br>Diving<br>Feeding<br>Fluking<br>Hunting<br>Hunting cooperative interspecific Yellowfin tuna (*Thunnus albacares*) with *Pseudorca crassidens*<br>Hunting cooperative interspecific (*Stenella frontalis*)<br>Leaping<br>Lining<br>Lunging<br>Play (with *Caretta caretta*)<br>Surfacing<br>Traveling<br>Traveling fast | 12 | 20 |

## Discussion

10 sighted species and 51 different behavioral patterns were observed in this 14-day survey. It emphasises the important role of the archipelago of the Azores for whale-watching as well as for whale research (Tobeña et al, 2016; Silva et al, 2003 und 2014). For practical treatment, the observed species are divided into two groups: huge whales, and Oceanic dolphins.

### Huge whales

In this study, two large whale species with quite similar body size were observed: *Physeter macrocephalus* (Odontoceti) and *Balaenoptera borealis* (Mysticetiy). Both species were often seen, but they vary in the number of different behavior patterns recorded. Sperm whale displayed 20 and Sei whale 10 behavioral patterns (Tab. 1). The result may be explained by their feeding ecology.

### Sei whale

The occurrence of this species in July/August and October is in contrary to Visser et al. (2011a). They reported that the abundance of Sei whales in the Azorean area is strongly related to the onset of the North Atlantic spring bloom. No sightings occurred during autumn. González Garcia et al. (2018) expressed another important issue related to the high dynamic nature of the marine environment in the Azores: a constantly changing environment created by the interaction between the dynamic oceanographic conditions and the wide range of sea depths. That provides a great variety of living conditions throughout the year. Probably prey availability was higher during this study than in other years.

Their behavioral budgets is largely composed of traveling and foraging. All baleen whales in the Azorean environment spent a substantial part of their time feeding. Foraging comprises over 40 % of the behavioral budget of Sei whales (Visser et al., 2011a). That is distinctly smaller than for the Sperm whale, but in this study, *Balaenoptera borealis* was observed traveling most of the time, which confirms Silva et al. (2014), Tobeña et al. (2016), and Pérez-Jorge et al. (2020) who suggested that the region of the Azores may be a transit area for Sei whales. During this study, *Balaenoptera borealis* sometimes interrupted traveling and feeding to slow locomotion and showed curiosity while approaching the boat.

### Sperm whale

For *Physeter macrocephalus* the Azores are both a breeding and a feeding area (Matthews et al., 2001; Magalhães et al, 2002). The waters around the Azorean Islands are one of the most important feeding ground for the species in the North Atlantic despite their low productivity (Oliveira, 2014). Especially the area south of Pico Island was one of the most prosperous Azorean whaling grounds. It is now one of the most important places for the recent whale-watching activities (Magalhães et al, 2002) with Sperm whale as a target species. *Physeter macrocephalus* has two main behavioral states, foraging and resting or socializing. They spend about 75% of its time foraging. They perform a series of long and deep dives to search and capture food (Magalhães et al, 2002). At the surface the foraging period are interspersed with a recovery time for female Sperm whales of about 10 min (Oliveira, 2014).

The Azores are frequented by females more often than by males. Bron et al. (2019) described regular sightings of very small groups or single individuals. (Silva et al (2014) reported groups of females accompanied by juveniles and calves observed foraging in the Azores every month. The forming of such groups is in agreement with this study. Aggregation and the prolonged recovery periods at the surface are explanations why twice as much behavior patterns have been observed for Sperm whale in comparison to Sei whale. During this study, Sperm whales represented a number of social and parental behavior patterns when resting, e.g. display, joining, lactating, play, and lob-tailing (Tab. 1) in compliance with Oliveira (2014), which *Balaenoptera borealis* did not show.

Mature males occasionally visit females to breed otherwise they are solitary (Magalhães et al, 2002). In this study, huge male Sperm whales were rarely observed. In all encounters the approach with the boat was executed very cautious. If a vessel gets too close they normally disappear, a behavior called “shallow dive”. Without showing any fluke they drop below the surface and may stay there for an indefinite time or they may slowly swim away to emerge far away. The observation on October 9^th^ was completely different. A huge male Sperm whale approached the boat on his own accord and showed his impressive dimension and a very rarely recorded imposing behavior.

### Oceanic dolphins

In this study, a few examples of interspecific cooperative hunting were registered (Tab. 1). Mixed-species feeding association of different Oceanic dolphin species together with large tunas and seabirds are a regular sighting, especially in the summer (Clua & Grosvalet, 2001). There are four species commonly involved, *Delphinus delphis, Tursiops truncatus, Stenella coeruleoalba*, and *Stenella frontalis* (Quérouil et al., 2008). Risso’s dolphin is rarely observed in association with one or the other species, which confirms this study. During all observed hunts, seabirds were watched as well.

Mixed-species aggregations were also observed. Aggregation occurs by chance, which distinguishes it from the functional association (Quérouil et al, 2008). In this study, two observations of aggregation were recorded. On July 24^th^ Sei whale and Spotted dolphins and on July 25^th^ Sperm whale with Common shortbeaked dolphin (Tab. 1).

### Shortbeaked common dolphin

In the Azores they use the area primarily for foraging and traveling (Cecchetti, 2017). Neumann (2001) described the group activity of *Delphinus delphis* with an overall proportion of 54.8% traveling, 20.5% milling, 17% feeding, 7,3% socializing, and 0.4% resting. The observation of this study confirms that Shortbeaked common dolphins spent most of their time traveling. Availability of prey is very likely governing the distribution.

*Delphinus delphis* was observed displaying a lower number of behavior patterns. One reason maybe the *Delphinus delphis* displacement by *Stenella frontalis* in summer and autumn. In this study, *Delphinus delphis* displayed bow riding, fluking, leaping, or snout-riding while traveling (Tab. 1). *Delphinus delphis* was observed in a mixed-species foraging association with *Stenella coeruleoalba* and *Calonectris diomedea borealis* on July 30^th^. Other activities can be assumed to become more frequent, only after nutritional needs have been satisfied (Neumann, 2001). They seldom approached the boat while they were feeding. In this study, resting was not observed, but sometimes aerial behavior. This confirms Cecchetti et al. (2018). They reported that *Delphinus delphis* occasionally exhibits conspicuous behaviors above water and tends to approach moving boats. This behaviors also aids their detection.

### Striped dolphin

In the region of Pico Island *Stenella coeruleoalba* usually avoids whale-watching boats. If a vessel approaches their school they often dash away at high speed (Carwardine, 2020). In this study, porpoising was the normal behavior to watch and *Stenella coeruleoalba* was easy to distinguish from other small dolphin species. So far there was only one known exception, when found in a mixed-species group. Than they endured a boat in their vicinity. In the morning on July 30^th^ *Stenella coeruleoalba* and *Delphinus delphis* were found forming an association for feeding. Therefore, eight different behavior patterns were observed, e.g. leaping, surfing, and traveling (Tab. 1).

### Atlantic spotted dolphin

In this study, Atlantic spotted dolphin represented 26 different behavior patterns, by far the most of all observed cetaceans (Tab. 1). A substantial number of energetic behavior was observed from *Stenella frontalis*, e.g. acrobatic leaps, somersaults, head- and tail-slapping, and tail-standing, which is in agreement to Carwardine (2020). Also different foraging behavior were recorded, e.g. bubble blowing, lining, pursuit, tail-slapping to stun fish, and interspecific cooperative hunting (Tab. 1).

Atlantic spotted dolphin have strong social bonds, especially between mother and calf. Segregation occurs by age and sex (Perrin, 2009), which this study confirms. On October 7^th^ an intraspecific association of several thousand individuals of Atlantic spotted dolphin, forming a superpod, occurred. The skipper reported that annual observations of associations like this are a not unusual in that period (personal communication). This sighting confirms the report of Silva et al (2014), which described an unusual number of sightings recorded by land-based encounters in October as well. Najarro (2020) reported an increase in average group size from arrival until departure in the Azores from 45 to 100 individuals.

### Common bottlenose dolphin

In this study, *<u>Tursiops truncatus</u>* was sighted twice as much as Shortbeaked common dolphin. The number of different behavior patterns recorded were 20, e.g. approach boat, bow riding, leaping, and shy-hopping, seven more in comparison to *Delphinus delphis*. The behavior resting was not observed, but approach boat and breaching (Tab. 1).

Three associations with other dolphins were recorded. The first one took place on August 1^st^ when Common bottlenose dolphin and Risso’s dolphin associated. Prior to this association a behavior of *Tursiops truncatus* was registered not mentioned in the literature so far. In the afternoon on August 1^st^ Common bottlenose dolphin was observed swimming very rapidly at the surface in a straight line for several hundred meters. This behavior was defined as “sprinting”. At the end of this line was a group of Risso’s dolphin. It rather seemed like they were the destination of *Tursiops truncatus*. The two groups united, approximately 10 individuals of each species associated. What followed seemed like aggressive behavior, because both species displayed fast chases, short dives, rapid surfacing, often fluking, frequent direction changes, producing a lot of splash water. Especially one behavior seemed very aggressive, when the individuals leaped upon another to force the lower animal under the surface, in the intention of drowning the other. Therefore, the behavior was defined as “drowning”.

In the morning of the next day both species were observed almost at the same location displaying the same behavior. Therefore, the assumption of the day before was rejected, that Risso’s dolphin tried to disperse Common bottlenose dolphin away from their habitat and that aggressive behavior was the prime reason. That matches the observation of the day before that only juvenile male *Grampus griseus* were engaged with *Tursiops truncatus*. During the observation that first day a few adult males arrived later on the spot. They did not interfere as expected, but remained close by. It was assumed that the juveniles, identified by their dark pigmentation, communicated with the adult males, distinguishable by their almost white coloration. After the arrival of the mature males the Risso’s dolphins outnumbered Common bottlenose dolphins. Since the adult males did not interfere in this situation the behavior of the primary group stayed unchanged when the boat departed.

The second observation lead to the assumption, that when both species met they showed social behavior. In this case they practiced aggressive behavior necessary in defense of this particular habitat to the other species. Both species differ in their feeding ecology that interspecific competition about nutrition can be ruled out. Pereira (2008b) described an interaction with Risso’s and Bottlenose dolphins in the Azores with chases, fast swimming, and leaps completely out of the water. The interaction ended with *Tursiops truncatus* leaving the area (Pereira, 2008b).

In this study, the two other associations of Bottlenose dolphin were observed in October. In both cases interspecific cooperative hunting was recorded. On October 7^th^ *Tursiops truncatus* and *Pseudorca crassidens* were registered hunting Yellowfin tuna (*Thunnus albacares*). Also intraspecific food sharing was observed within each species during the hunt. On October 8^th^ Common bottlenose dolphin and Atlantic spotted dolphin were observed with interspecific cooperative hunting for small fish. Also on October 7^th^ *Tursiops truncatus* playing with Loggerhead sea turtle (*Caretta caretta*) was recorded.

### Short-finned pilot whale

This study revealed eight different behaviour (Tab. 1). Because of the mainly nocturnal feeding behavior, this species spend much of the day logging at the surface (Cawardine, 2020). Since they are seldom very active during daytime the behavior mostly displayed are slow traveling, resting, and sometimes fluke-slapping (Sajikumar et al, 2014). In this study, socializing was not observed, although *Globicephala macorhynchus* has a highly social nature. Therefore, the finding may appear unusual. Jensen et al. (2011) also never observed social behavior in their study of Short-finned pilot whales. The result of this study confirms Shane (1995) who found Short-finned pilot whales off Santa Catalina Island, California, traveling 73 % of the time. In this study, diving and traveling were the most observed behavioral patterns. Traveling and diving behavior is usually classified as an avoidance response.

### Risso’s dolphin

In this study, *Grampus griseus* represented a high number of behavioral patterns (20), just as *Tursiops truncatus* (Tab. 1). The registration of a diversity of energetic behaviors such as flipper- and tail-slapping, lateral swimming, and leaping are in agreement with Pereira (2008b). It also confirms Hartman et al (2009) that Risso’s dolphins occur in relatively small groups characterized by a high degree of synchrony and calm-surfacing. Synchronous behavior is often displayed by males.

*Grampus griseus* resting was not observed in this survey. This might be explained by Visser et al. (2011b) findings, that Risso’s dolphins alter their daily resting pattern in response to whale-watching in the Azores. In the low season *Grampus griseus* groups rested mainly in the morning and in the afternoon. During the high season, when more whale-watching vessels are out, they rested less and did so mainly at noon. At that time the number of vessels was lowest. In this study, the boat tours were executed in the described manner. The observations of this study confirms Pereira (2008b), who reported for *Grampus griseus* diurnal activities mostly traveling (77%), socializing (13%), feeding (5%), and resting (4%) in the Azores. And also Shane (1995), who found Risso’s dolphins off Santa Catalina Island, California, traveling 84 % and resting 1 % of the time.

Pereira (2008b) stated two species groups and interactions with six cetacean species in the Azores. In this study, association with *Tursiops truncatus* was reported and discussed in a previous chapter. In comparison to this study, with 12 sightings of Risso’s dolphins, Pereira (2008b) collected data on 107 sightings, which justifies the smaller number of interactions. Pereira (2008b) also describes harassment behaviors with Globicephala spp. and *Physeter macrocephalus* suggesting competitive interference. Possibly the agonistic interaction with Common bottlenose dolphin discussed the previous chapter is related to this observation. Probably in the context of practising agonistic behavior with a species of the same size, and to a lesser extend an interspecific competition.

### False killer whale

*Pseudorca crassidens* is an exuberant and fast-swimming cetacean and often leaps clear out of the water (Carwardine, 2020), which this study confirms. With four observations in October, this species was rarely sighted. Bow riding, breaching, lining, lunging, spy-hopping, tail-slapping, with 22 recorded behavior patterns the False killer whale demonstrated the second highest number of all observed cetaceans (Tab. 1).

*Pseudorca crassidens*, also rarely observed, associate with a number of other odontocete species, in particular with *Tursiops truncatus* (Zaeschmar et al, 2012). This study confirms this with the recording of an interspecific cooperative hunting. False killer whale and Common bottlenose dolphin were watched foraging on Yellowfin tuna (*Thunnus albacares*). To date there is no known record of this behavior observed in the Azores. Both species were interspersed into mixed-species subgroups over an area of approximately 8 km^2^. The foraging was indicated by leaps, asynchronous dives and the association of sea birds in agreement to Zaeschmar et al (2012). Sometimes False killer whales were observed carrying caught Yellowfin tuna in their mouth.

This observation led to another one, the intraspecific food sharing within the species. Food sharing is defined as the unresisted transfer of food (Jaeggi & Schaik, 2011). They define possession as being in physical contact with the food. That excludes transfers without clear possession, such as collecting scraps from the vicinity of a feeding individual, e.g. sea birds snatching particles in the area of foraging dolphins. Therefore, food sharing is the voluntarily relinquish of food from a possessor to the recipient. It is the benefit of the receiver (Jaeggi & Schaik, 2011). The passing of large pieces of tuna from one False killer whale to another one was observed in agreement with Jaeggi & Schaik (2011). Wright et al (2016) reported food sharing in *Orcinus orca* was nonreciprocal and maternal relatedness was a significant predictor of the frequency of prey sharing. In this study, it was not possible to determine if the sharing took place with related offspring or not.

## Conclusions

Data used in this research were collected as a by-product of whale-watching, which was the primary purpose. The normal tourism activity favour the observation of different species (Pereira, 2008b). This method is subject to various limitations. The study area may not be equally covered. Therefore, coincidence may play an important role. Findings from this research may not be representative for the entire research area. The boat surveys during the excursions were not designed for the purpose to estimating cetacean abundance (Bron et al, 2019; Silva et al, 2014). Another disadvantage of this method may be that it provides information on the behavior of potentially disturbed whales (Magalhães et al, 2002).

Nevertheless, the activity of whale-watching offer a source of valuable data. It can be used for sighting surveys, e.g. to obtain an estimate of the size of the population in the study area. It represents a cost-effective method to collect data on the cetaceans. Otherwise the information may be inaccessible, e.g. for rare species or incidental sightings. This study obtained data that may be helpful to a better understanding of the role of sustainable whale-watching.

Whale-watching tours are often offered year-round and the data collected may be a potential tool for detecting long term changes (Bron et al, 2019; Silva et al, 2014). The International Whaling Commission (IWC) describes in its whale-watching handbook the common research methods used to study whales and dolphins. They recommend a few of these methods to be easily combined with whale-watching. One of these are boat surveys to study distribution and habitat use (IWC, 2020). The knowledge obtained here in this study could be used for policies aimed at effectively protecting cetaceans and their habitats or in the development of management plans for specific areas, e.g. the definition of a load capacity for whale-watching activities (Silva et al, 2003; Tobeña et al, 2016).

## Acknowledgments

Sincere thanks are given to the staff of Espaço Talassa for their friendly support in all matters of the preparation and execution of the excursions. We acknowledge the help of the students throughout this study, their commitment and enthusiasm for the protection of whales.

